# Transcriptome-wide association and prediction for carotenoids and tocochromanols in fresh sweet corn kernels

**DOI:** 10.1101/2021.09.24.461734

**Authors:** Jenna Hershberger, Ryokei Tanaka, Joshua C. Wood, Nicholas Kaczmar, Di Wu, John P. Hamilton, Dean DellaPenna, C. Robin Buell, Michael A. Gore

## Abstract

Sweet corn is consistently one of the most highly consumed vegetables in the U.S., providing a valuable opportunity to increase nutrient intake through biofortification. Significant variation for carotenoid (provitamin A, lutein, zeaxanthin) and tocochromanol (vitamin E, antioxidants) levels is present in temperate sweet corn germplasm, yet previous genome-wide association studies (GWAS) of these traits have been limited by low statistical power and mapping resolution. Here, we employed a high-quality transcriptomic dataset collected from fresh sweet corn kernels to conduct transcriptome-wide association studies (TWAS) and transcriptome prediction studies for 39 carotenoid and tocochromanol traits. In agreement with previous GWAS findings, TWAS detected significant associations for four causal genes, *β-carotene hydroxylase (crtRB1)*, *lycopene epsilon cyclase* (*lcyE*), *γ-tocopherol methyltransferase* (*vte4*), and *homogentisate geranylgeranyltransferase* (*hggt1*) on a transcriptome-wide level. Pathway-level analysis revealed additional associations for *deoxy-xylulose synthase2* (*dxs2*), *diphosphocytidyl methyl erythritol synthase2* (*dmes2*), *cytidine methyl kinase1* (*cmk1*), and *geranylgeranyl hydrogenase1* (*ggh1*), of which, *dmes2, cmk1*, and *ggh1* have not previously been identified through maize association studies. Evaluation of prediction models incorporating genome-wide markers and transcriptome-wide abundances revealed a trait-dependent benefit to the inclusion of both genomic and transcriptomic data over solely genomic data, but both transcriptome- and genome-wide datasets outperformed *a priori* candidate gene-targeted prediction models for most traits. Altogether, this study represents an important step towards understanding the role of regulatory variation in the accumulation of vitamins in fresh sweet corn kernels.

**Core Ideas:**

1. Transcriptomic data aid the study of vitamin levels in fresh sweet corn kernels.
2. *crtRB1, lcyE, dxs2, dmes2*, and *cmk1* were associated with carotenoid traits.
3. *vte4, hggt1*, and *ggh1* were associated with tocochromanol traits.
4. Transcriptomic data boosted predictive ability over genomic data alone for some traits.
5. Joint transcriptome- and genome-wide models achieved the highest predictive abilities.

## INTRODUCTION

Micronutrient deficiencies affect more than two billion people across the globe (FAO 2020), with most cases occurring in developing countries. Although clinical deficiency is less common in the U.S., nutrient intake below the estimated average requirement is still prevalent in vulnerable sectors of the population (Wallace et al. 2014; Eggersdorfer et al. 2018). Essential micronutrients such as vitamins A and E are necessary for normal biological functioning and suboptimal levels of these nutrients can contribute to health complications. Inadequate vitamin E intake has been linked to increased risk of cardiovascular diseases (Knekt et al. 1994; Kushi et al. 1996), and lutein and zeaxanthin, two non-provitamin A dietary carotenoids, have been associated with reduced risk of age-related macular degeneration (Wu et al. 2015). The primary sources of vitamins A and E are dietary carotenoids and tocochromanols, respectively, but bioavailability of these nutrients varies depending on the food source and preparation (Yeum and Russell 2002;Tanumihardjo et al. 2010; Borel et al. 2013).

Several strategies for improvement of micronutrient intake have been employed in the U.S. including industrial fortification of commonly-consumed foods, multivitamin and mineral supplementation, nutritional education for promotion of dietary diversification, and biofortification (Allen et al. 2006). In terms of long-term impact, investment in biofortification has the potential to provide the most lasting benefit, requiring no additional funding beyond upfront research and development costs. The biofortification of staple crops through both conventional and molecular breeding techniques has been successful in increasing the nutritional content of new plant varieties through initiatives such as HarvestPlus (Bouis and Saltzman 2017;Díaz-Gómez et al. 2017; Hirschi 2020; Bhullar and Gruissem 2013). Although few have focused on U.S. impact, some of these efforts have targeted nutrients lacking in many U.S. diets such as carotenoids (Low et al. 2017) and tocochromanols (Che et al. 2016), thus serving as examples for biofortification efforts in the U.S. Sweet corn is routinely ranked amongst the most highly consumed vegetables in the U.S. (USDA ERS 2020), and adequate heritable variation exists for improvement of carotenoid and tocochromanol traits in fresh sweet corn kernels (Baseggio et al.2019; Baseggio et al. 2020; Xiao et al. 2020). Paired with the many genomic resources that can be leveraged from the greater maize germplasm pool, these attributes position sweet corn as an ideal target for biofortification.

Biofortification is enabled by our growing knowledge of the biosynthetic pathways and variants controlling phenotypic diversity within a population, especially when powered by genomics-assisted selection. While carotenoid and tocochromanol traits have been well-characterized in mature maize grain through linkage analysis and genome-wide association studies (GWAS) (Harjes et al. 2008; Yan et al. 2010; Li et al. 2012; Lipka et al. 2013; Owens et al. 2014; Diepenbrock et al. 2017; Diepenbrock et al. 2021), sweet corn is a distinct subpopulation of maize with allele frequencies that differ from those of the broader maize germplasm base (Romay et al. 2013), potentially limiting the transferability of these findings. Genetic dissection of carotenoid and tocochromanol accumulation in fresh sweet corn kernels has revealed relatively fewer insights into the genetic control of these traits, with outcomes limited by low mapping resolution and confounding associations of some endosperm-specific traits with *sugary1* (*su1*) and *shrunken2* (*sh2*), kernel endosperm starch mutations characteristic of sweet corn (Baseggio et al. 2019; Baseggio et al. 2020). The use of gene expression data, offering gene-level resolution and insight into regulatory variation, has the potential to help overcome these obstacles. Transcriptome-wide association studies (TWAS), which assess the association between gene expression and terminal phenotypes (Hirsch et al. 2014; Lin et al. 2017; Pasaniuc and Price 2017; Li et al. 2021), have successfully identified associations previously found in GWAS and have highlighted new and promising associations for mature grain carotenoids and tocochromanols (Kremling et al. 2019).

In general, genomic selection leverages dense genomic marker data for generating genomic estimated breeding values to inform selection decisions, but its success is dependent on the predictive abilities of the underlying statistical models. The models commonly employed for genomic prediction (GP), such as GBLUP, assume numerous loci with small effects (Meuwissen et al. 2001). However, this assumption is in contrast with the genetic architecture of carotenoid and tocochromanol levels in physiologically mature maize grain and fresh sweet corn kernels, which consists of few genes with large effects (Lipka et al. 2013; Owens et al. 2014;Diepenbrock et al. 2017; Diepenbrock et al. 2021; Baseggio et al. 2019; Baseggio et al. 2020). This suggests that an informed, unequal weighting of markers according to phenotypic variation explained by causal genes would lead to more accurate predictions by GP models, a hypothesis supported by studies in multiple biological systems (van ’t Veer et al. 2002; Edwards et al. 2016;Fang et al. 2017).

Several approaches have been used to incorporate biological information into GP models for maize kernel tocochromanol and carotenoid traits, including quantitative trait loci (QTL) identified via linkage analysis. Moderately high predictive abilities have been achieved when utilizing both whole-genome prediction and smaller, carotenoid QTL-targeted marker sets in mature kernels of a maize association panel (Owens et al. 2014). In contrast, predictive abilities for many fresh kernel carotenoid and tocochromanol traits in a sweet corn association panel have been lower for QTL-targeted as compared to whole-genome marker sets when using joint-linkage (JL)-QTL identified in the maize nested association mapping (NAM) population (Baseggio et al. 2019; Baseggio et al. 2020). As only a fraction of these QTL were identified as significant associations in this same sweet corn panel (Baseggio et al. 2019; Baseggio et al.2020), it is possible that key causal variants were insufficiently represented in the targeted marker set.

The incorporation of gene expression data into GP models has the potential to increase predictive ability in sweet corn nutritional quality traits by providing additional information, such as regulatory variation, that may not be captured by genomic variation alone. Any improvements in the identification of causal genes through TWAS would also help to customize targeted GP models to better fit the needs of sweet corn. Models that leverage expression data have been shown to increase prediction abilities over genomic marker-based models, but increases are not consistent for all tested traits (Guo et al. 2016; Schrag et al. 2018; Azodi et al.2020). To our knowledge, these models are yet untested in sweet corn. We hypothesize that the sum of these advantages may allow us to address the challenges faced in previous GWAS and GP models with nutritional traits in sweet corn, resulting in a more deeply resolved genetic architecture and increased predictive ability.

The objectives of this study were to integrate gene expression data with existing genomic and phenotypic datasets to 1) better characterize the genetic architecture of carotenoid and tocochromanol concentrations in fresh sweet corn kernels, and 2) evaluate the potential of transcriptomic data to improve predictive abilities for genomic selection in a sweet corn biofortification breeding program.

## MATERIALS AND METHODS

### Germplasm and experimental design

A sweet corn association panel of 382 inbred lines representative of U.S. temperate breeding program genetic diversity were selected from the panel studied by Baseggio et al. (2019; 2020) based on the availability of single-nucleotide polymorphism (SNP) marker genotype and fresh kernel carotenoid and tocochromanol phenotype data in those studies. Inbred lines with *sugary1* (*su1*), *shrunken2* (*sh2*), and both *sugary1* and *shrunken2* (*su1sh2*) starch-deficient endosperm mutations were included, while the less common *aeduwx* or *bt2* mutations and lines with missing phenotypic data for both studies were excluded from this study.

The 382 lines were planted in an augmented incomplete block design as previously described (Baseggio et al. 2019). Briefly, three sets organized by plant height were used to reduce shading effects on shorter plants, with each set consisting of incomplete blocks of 20 single-row experimental plots. Each incomplete block was augmented with two of four height-specific checks (We05407 and W5579, W5579 and Ia5125, or Ia5125 and IL125b). Twelve seeds were planted in each plot. The experiment was planted in field N (Lima silt loam soil) at Cornell University’s Musgrave Research Farm in Aurora, NY on June 7, 2019. Plots were 3.05 m long with inter-row spacing of 0.76 m and a 0.91 m alley at the end of each plot.

A single ear per plot was harvested at approximately 400 growing-degree days (GDD) after self-pollination and flash-frozen with liquid nitrogen in the field. When possible, samples were not collected from the end plants of each plot. Ears were kept on dry ice during transportation to the fieldhouse laboratory, where the middle third of each ear was shelled and transferred into a sample cup. These samples were also kept on dry ice during transportation to a −80°C freezer. In total, 373 experimental and 69 check kernel samples were collected.

### RNA isolation and 3′ mRNA sequencing

Seven frozen kernels from each sample cup were ground in liquid nitrogen using an IKA grinder (IKA Works, Inc., Staufen, Germany). RNA was extracted from approximately 100 mg of ground tissue from each sample with hot borate and lithium chloride (Wan and Wilkins 1994). Extracted samples were DNAse treated with the Ambion Turbo DNA-free Kit (Thermo Fisher Scientific, Waltham, MA). The extraction process was repeated for degraded samples as identified with agarose gel electrophoresis. After isolation, RNA was quantified using a BioTek Epoch 2 microplate spectrophotometer (BioTek Instruments, Inc., Winooski, VT) and diluted to a concentration of 200-300 ng/μl. Diluted samples were randomly assigned to five 96-well plates. Two positive control wells containing pooled RNA from one of the field check inbred lines (W5579) and two negative control wells (one blank and one water) were included per plate. The five plates were submitted to the Cornell Biotechnology Resource Center (Cornell University, Ithaca, NY, USA) where libraries were constructed using the Lexogen QuantSeq 3′ mRNA-Seq Library Kit FWD (Lexogen, Greenland, NH). Constructed libraries were sequenced on an Illumina NextSeq 500 (Illumina, San Diego, CA) with a single plate per sequencing lane.

### Expression abundance estimation

The 3′ mRNAseq reads were cleaned in accordance with Lexogen recommendations (https://www.lexogen.com/wp-content/uploads/2020/04/015UG009V0252_QuantSeq_Illumina_2020-04-03.pdf) by completing two rounds of Cutadapt (v2.10) (Martin 2011). To summarize, round one trimmed Illumina adapters and round two trimmed the first 12 bases and polyA tails. Alignments were generated to the B73 v4 genome (Jiao et al. 2017) and the Ia453 genome (Hu et al. 2021) using HISAT2 (v2.2.1) (Kim et al. 2019) with the following parameters; --max-intronlen 5000, --dta-cufflinks, and --rna-strandness F. The htseq-count function from HTSeq (v0.12.4) (Anders et al. 2015) was used to generate counts with the following parameters; --order=pos, --stranded=yes, --minaqual=10, --type=gene, and --mode=union. This was done in a genome-dependent manner using the B73 v4.59 or Ia453 v1.0 annotations. The count data were normalized using the *rlog* function of DESeq2 (Love et al. 2014). Genes with a normalized count of less than or equal to zero for all samples were filtered out of the final count matrix. The normalized data were then filtered by removing samples that had high levels of contaminants (12 samples) that were introduced by the sequencing facility and further filtering out samples with less than 250,000 reads (3 samples). This resulted in a high-quality dataset of 433 samples.

Additional stringent filtering steps were performed to further prepare the dataset for downstream analyses. A total of 77 positive control and check samples were removed, resulting in 356 experimental samples. All genes with 50% or more samples having normalized transcript abundance values of zero were removed from the analysis. Extreme observations exceeding 100 median absolute deviations from the median for a given gene were removed from the analysis following the method of Davies and Gathers (1993), resulting in 0.01% of total observations removed in this step. These observations were imputed using the median normalized count value for that gene, as no missing values were permitted in the downstream calculation of probabilistic estimation of expression residuals (PEER) factors. Finally, genes with greater than 10% of observations above the median absolute deviation threshold were removed from the analysis. Filtering steps and metrics for each reference genome alignment are available in Supplemental Table S1.

### Statistical analysis of transcriptomic data

Best linear unbiased estimators (BLUEs) were calculated using the normalized transcript abundances for all genes passing quality control. The following mixed model was fitted to each gene with ASReml-R version 3.0 (Gilmour et al. 2009):

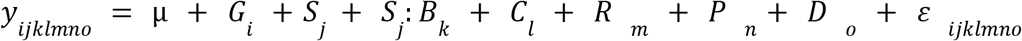

Where y_ijklmno_ is an individual normalized transcript abundance observation. μ is the overall mean, *G_i_* is the fixed effect for genotype *i*, *S_j_* is the random effect for set *j, B_k_* is the random effect for block *k* in set *j, C_l_* is the random effect for plot grid column *l*, *R_m_* is the random effect for plot grid row *m*, *P_n_* is the random effect for RNA sample 96-well plate *n*, *D_o_* is the random effect for RNA extraction date *o*, a categorical variable with two levels representing extraction dates before and after laboratory closures due to COVID-19, and *ε_ijklmno_* is the residual effect. Model fitting did not converge for a subset of genes for each reference genome, so transcript abundance BLUE values of 18,765 (B73) and 18,477 (Ia453) genes were obtained for 355 inbred lines (Supplemental Table S1).

PEER (Stegle et al. 2010), a Bayesian factor analysis which removes hidden factors (*e.g*., experimental noise) from the transcript abundance data, was used to prepare the expression profile for downstream analysis. First, an optimal number of factors was visually identified by finding the “elbow” of the diagnosis plot of the factor relevance (so-called scree plot) up to 25 factors (Supplemental Table S2). This PEER calculation was applied after removing genotypes that were not included in the phenotype dataset (four genotypes were removed for tocochromanol data; 72 genotypes were removed for carotenoid data). Finally, a linear model with grand mean was fitted on the PEER residuals for each gene and Studentized deleted residuals (Neter et al. 1996) were used for outlier identification given a Bonferroni correction (α = 0.05).

### Existing phenotypic data

The sweet corn association panel was previously evaluated for fresh kernel tocochromanol and carotenoid traits during the 2014 and 2015 field seasons at Cornell University’s Musgrave Research Farm in Aurora, NY as previously described (Baseggio et al.2019; Baseggio et al. 2020). BLUPs in μg g^-1^ for these 20 tocochromanol traits [δ-tocotrienol, δT3; γ-tocotrienol, γT3; α-tocotrienol, αT3; δ-tocopherol, δT; γ-tocopherol, γT; α-tocopherol, αT; total tocotrienols, total T3; total tocopherols, total T; total tocochromanols, total T3 + T; αT3/γT3; αT/γT; δT3/αT3; δT/αT; δT3/γT3; δT/γT; γT3/(γT3 + αT3); γT/(γT + αT); δT3/(γT3 + αT3); δT/(γT + αT); total T/total T3] from Baseggio et al. (2019) and 19 carotenoid traits [antheraxanthin; β-carotene; β-cryptoxanthin; lutein; violaxanthin; zeaxanthin; zeinoxanthin; other carotenes (lycopene, α-carotene, δ-carotene, and other unidentified carotenes); zeinoxanthin/lutein; β-cryptoxanthin/zeaxanthin; β-carotene/β-cryptoxanthin; β-carotene/(β-cryptoxanthin+zeaxanthin); α-xanthophylls (sum of lutein and zeinoxanthin); β-xanthophylls (sum of antheraxanthin, β-cryptoxanthin, violaxanthin, and zeaxanthin); β-xanthophylls/α-xanthophylls; total carotenes (sum of β-carotene and other carotenes); total xanthophylls (sum of α- and β-xanthophylls); total carotenes/total xanthophylls; total carotenoids (sum of the seven carotenoid compounds and other carotenes)] from Baseggio et al. (2020) were used for association and prediction analyses in this study.

### Transcriptome-wide association studies

TWAS was performed for each of the 39 fresh kernel tocochromanol and carotenoid traits using transcripts aligned to the B73 v4 and Ia453 reference genomes. An additional TWAS was conducted for kernel type mutations (*sh2* or *su1*) with both reference genome alignments, but because only 17 lines contained both *sh2* and *su1* kernel type mutations, lines including both mutations were excluded from the analysis of this trait. Prior to conducting TWAS, a total of 44 models were compared for each tocochromanol and carotenoid trait: with or without kinship (*n* = 2 options), with or without kernel mutant type (*n* = 2 options), and different numbers of principal components (PCs) (*n* = 11 options; 0-10 PCs). Kernel mutant type was not included as a fixed effect in models for the kernel type trait, thus a total of 22 models were evaluated for this trait. For models including kinship, a random genotypic effect was also included in the model. If included in the optimal model, kernel mutant type and PCs were modeled as fixed effects. The Bayesian information criterion (BIC) was used to select the optimal model for each trait (Supplemental Table S3). Kinship matrices were calculated with SNPs having a minor allele frequency > 0.05 obtained from the 10,773 SNP dataset from Baseggio et al. (2021) according to VanRaden’s method 1 (VanRaden 2008) implemented in GAPIT version 3 (Lipka et al. 2012). TWAS was implemented using the optimal model for each trait with the GWAS() function in the rrBLUP R package (Endelman 2011). Resulting raw *P*-values were adjusted to control the false discovery rate (FDR) using the p.adjust() function in base R (Benjamini and Hochberg 1995; R Core Team 2018). Significant associations were identified as those passing a 5% FDR threshold.

### Pathway-level analysis

A pathway-level analysis considering only *a priori* candidate pathway genes in the well-maintained and annotated B73 reference genome was employed to increase the probability of identifying weaker-effect alleles contributing to carotenoid and tocochromanol phenotypic variation (Lipka et al. 2013; Owens et al. 2014). A list of *a priori* candidate genes involved in accumulation (carotenoids and tocochromanols) and retention (carotenoids) was compiled from previous studies of carotenoid and tocochromanol levels in maize kernels (Diepenbrock et al.2017; Diepenbrock et al. 2021) and further updated for tocochromanols in our study. The common support intervals of JL-QTL from the U.S. maize NAM panel were uplifted from B73 v2 to B73 v4 according to the methods described by Wu et al. (2021). These common support intervals and *a priori* candidate genes were matched with B73 v4-aligned TWAS results to facilitate the biological interpretation of findings (Supplemental Table S4). Using subsets including only the genes identified as *a priori* candidates for each set of traits, pathway-level FDR adjustments were performed on the raw (unadjusted) TWAS *P*-values for both the tocochromanol and carotenoid trait sets with the p.adjust() function in base R. A threshold of 5% FDR was used to declare significant associations.

### Genome- and transcriptome-wide prediction

Genomic relationship matrices (GRMs) were calculated according to VanRaden’s method 1 in GAPIT (VanRaden 2008; Lipka et al. 2012) using a set of 147,762 GBS SNPs from Baseggio et al. (2021) uplifted to B73 v4 (Jiao et al. 2017) with minor allele frequency > 0.05. Transcriptomic relationship matrices (TRMs) were calculated for each reference genome by scaling the cross-product of the centered and standardized expression-BLUE matrix by the number of transcripts (VanRaden 2008; Morota and Gianola 2014).

We built 11 unique models using six relationship matrices: five with a single relationship matrix (GRM, TRM.B73, TRM.Ia453, TRM.cand, and TRM.non.cand) and six with multiple relationship matrices (GRM+TRM.B73, GRM+TRM.Ia453, GRM+TRM.cand, GRM+TRM.non.cand, TRM.both, and GRM+TRM.both) (Table 1). Relationship matrices were additively included for models with more than one relationship matrix (*i.e*., multi-kernel prediction). All models were tested with and without kernel mutant type included as a covariate. When included, kernel mutant type was modeled as a fixed categorical variable with three levels (*sh2, su1*, and *su1sh2*). All models were implemented in the BGLR package in R (Pérez and De Los Campos 2014), with the number of iterations for MCMC set to 12,000 with 8,000 samples for the burn-in period. All other parameters were set to the default values. Five-fold cross validation was repeated 10 times to evaluate the predictive ability (Pearson’s correlation between predicted and observed values) for each phenotype. Folds were stratified by endosperm mutant type (*sh2*, *su1*, and *su1sh2*) to represent population genotype frequencies and were held consistent across all models.

**Table 1.**
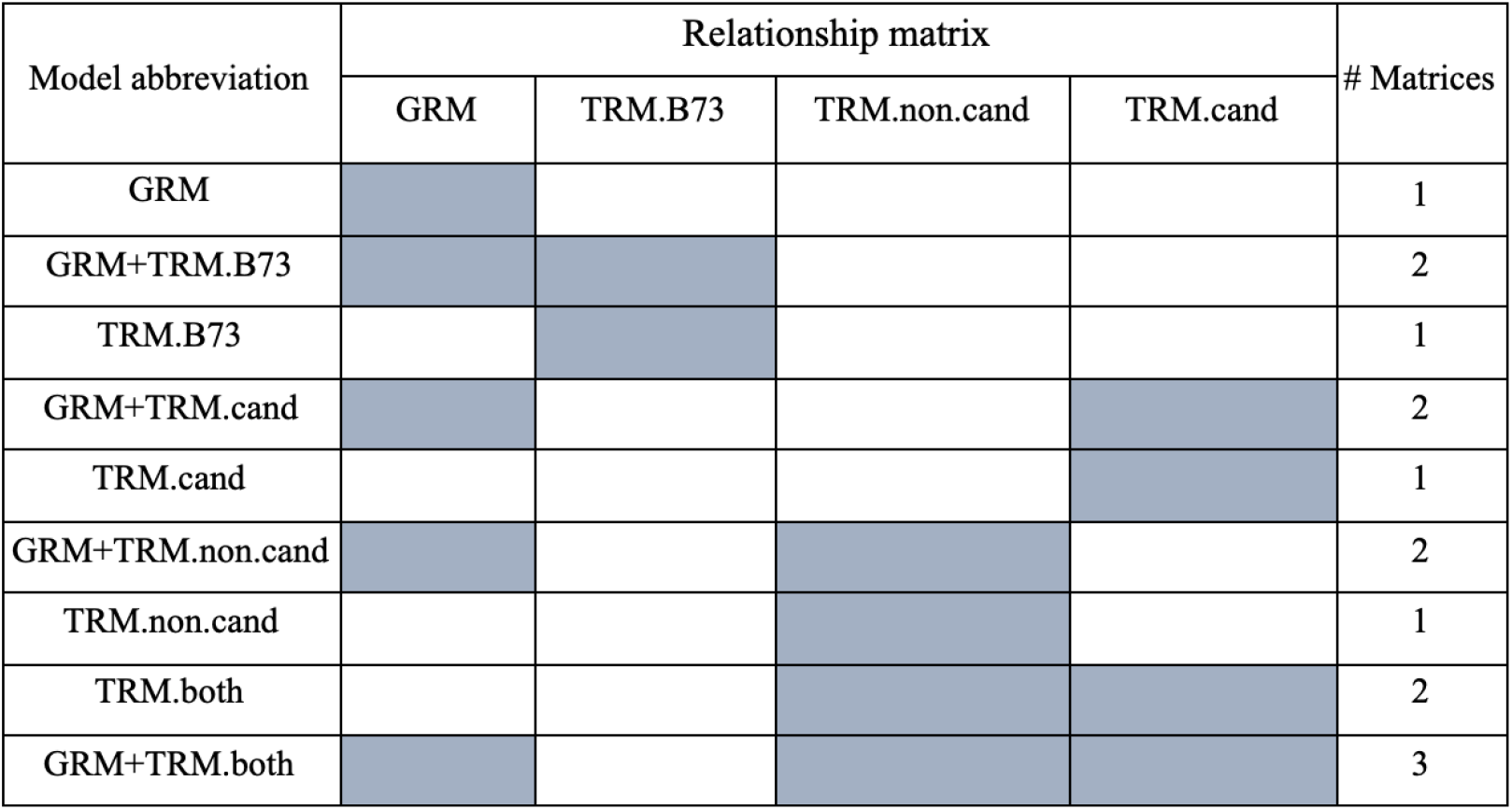
Summary of the models evaluated in this study GRM is the genomic relationship matrix, TRM.B73 is the transcriptomic relationship matrix using all available genes, TRM.cand is the transcriptomic relationship matrix incorporating only *a priori* candidate genes, and TRM.non.cand is the transcriptomic relationship matrix using only genes categorized as not *a priori* candidates. A list of included or excluded candidate genes is available in Supplemental Table S4. Models using transcriptomic data aligned to the Ia453 reference genome (TRM.Ia453) follow the same form as those using TRM.B73 shown here, but the transcriptomic data used to calculate the TRM.cand and TRM.non.cand matrices were aligned to the B73 v4 reference genome only.

## RESULTS

### Kernel type TWAS

As a proof of concept, we performed TWAS for kernel mutation type in a diverse panel of sweet corn inbred lines representative of temperate U.S. germplasm. As we defined it, this trait is caused by mutations in two genes known to be involved in endosperm starch biosynthesis, *sh2* and *su1*, and was represented in a binary manner in this analysis. Due to structural differences between the B73 and Ia453 genomes at the *sh2* locus (Hu et al. 2021), transcripts from fresh kernels were aligned to both genomes to enable comparison between the two. Ia453 has the *sh2-reference* mutant allele (*sh2-R*), which consists of two genes that originated from a single copy of the functional *sh2* allele as the result of structural rearrangement, while B73 has the functional *sh2* allele.

Strong associations were observed between kernel mutant type and transcript abundance of *sh2* using both the B73 (Zm00001d044129; *P*-value 3.20 × 10^−26^) and Ia453 (Zm00045a021195; *P*-value 1.31 × 10^−26^) reference genomes at a transcriptome-wide 5% FDR. Transcript abundance of the second *sh2* gene in the Ia453 reference genome, Zm00045a021196, was not significantly associated with the kernel mutant type trait in our panel (*P*-value 6.73 × 10^−2^). Despite a presumed causal relationship between mutations in the *su1* gene and the resulting kernel phenotype, a significant association was not identified with transcript abundance of this gene in our panel with either reference alignment (Figure 1). Supporting these results, there is little variation overall for *su1* expression regardless of visual and marker-based kernel endosperm mutation classification, while a relatively large range in transcript abundance is observed at *sh2* based on these same groupings (Figure 2).

**Figure 1.**
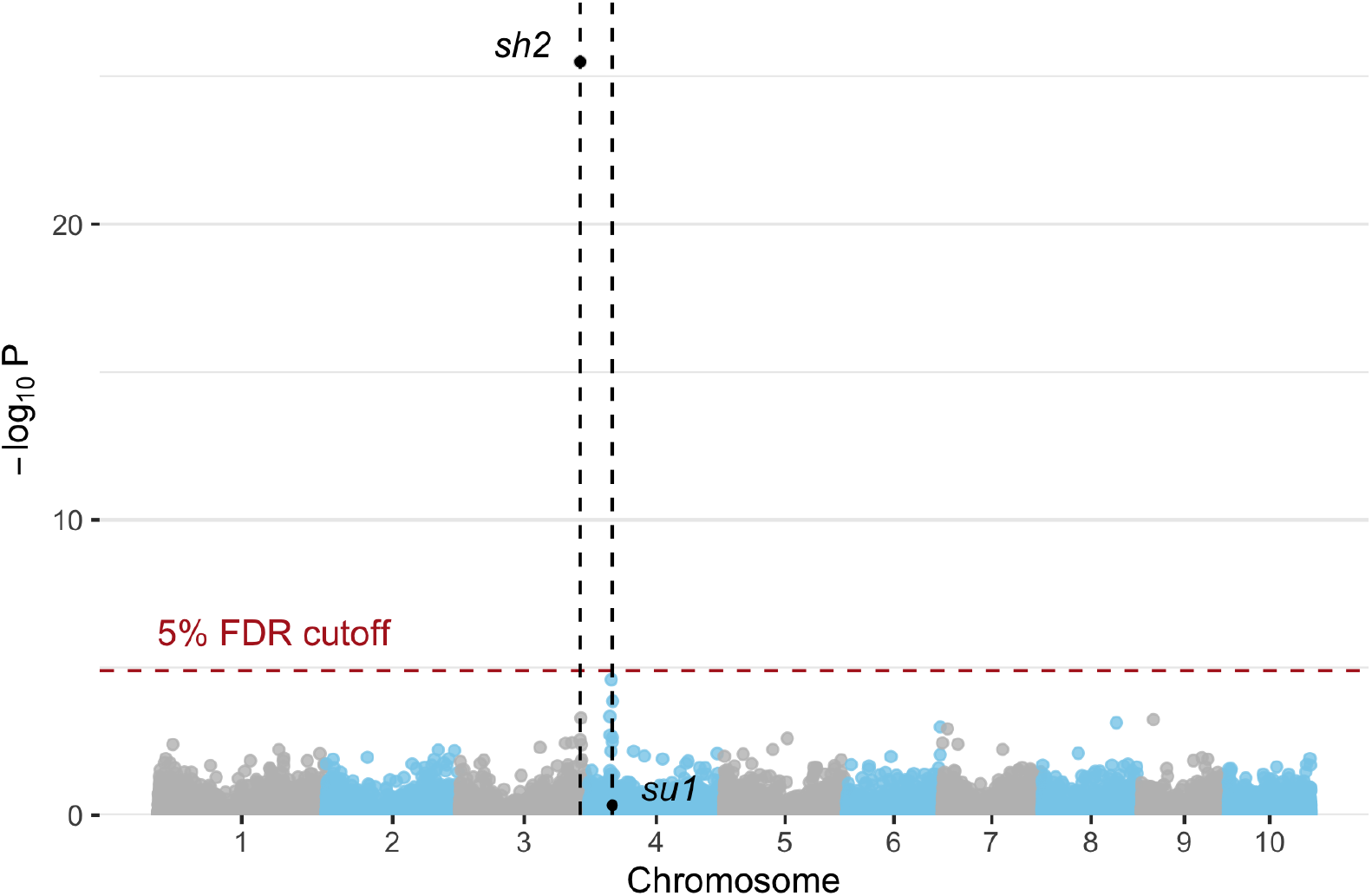
Manhattan plot representing the -log_10_ *P*-values of the transcriptome-wide association study of endosperm mutation type using the B73 v4 reference genome. Relative positions of *sh2* (*shrunken2*; Zm00001d044129) and *su1* (*sugary1*; Zm00001d049753) are indicated by black dots and vertical lines.

**Figure 2.**
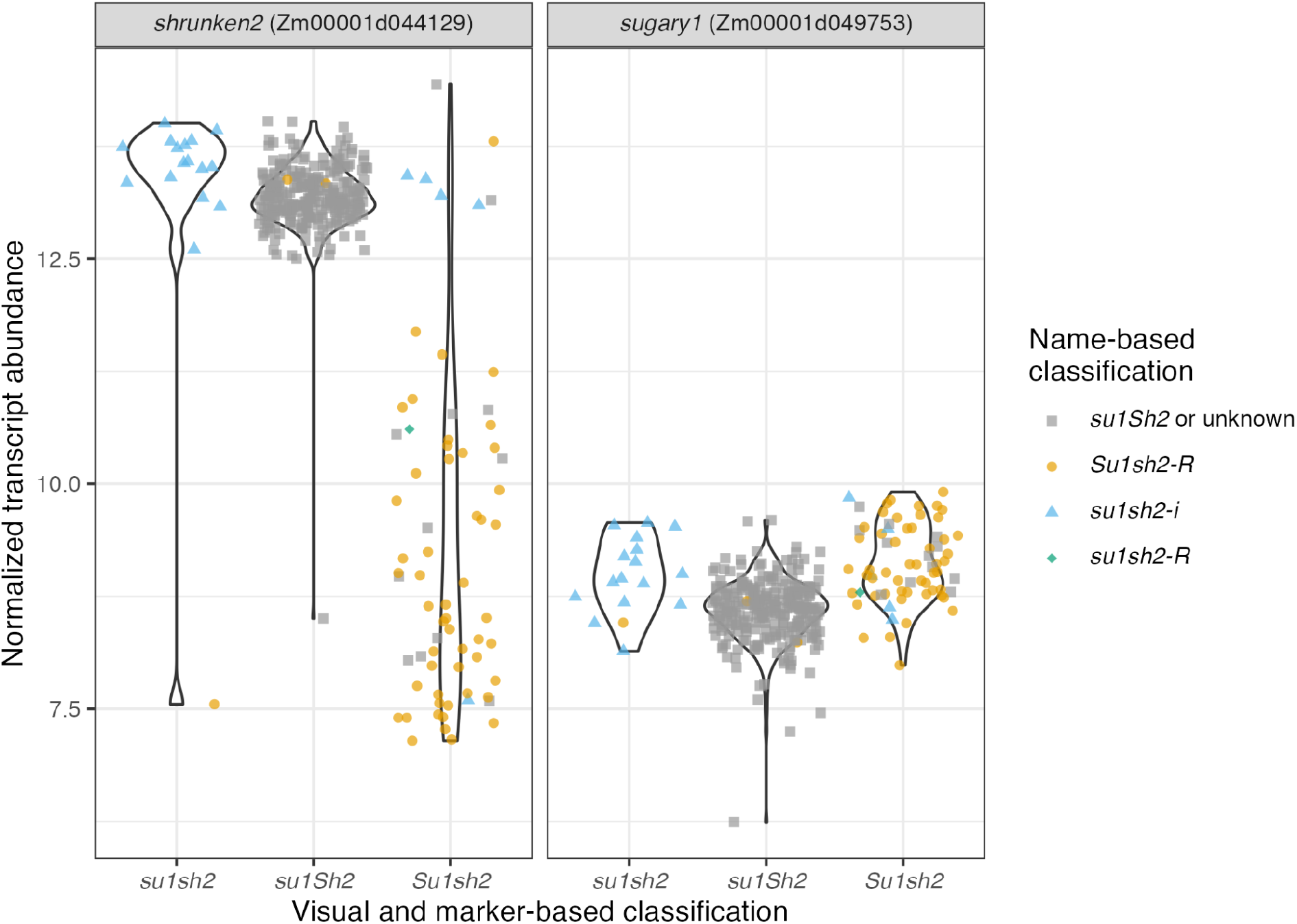
B73-aligned normalized transcript abundance of fresh sweet corn kernels by kernel endosperm mutation type assignment for *sh2* (*shrunken2*; Zm00001d044129) and *su1* (*sugary1*; Zm00001d049753). Visual and marker-based endosperm classifications on the x-axis are from Baseggio et al. (2019; 2020), while name-based classifications (color and shape of data points) are based on inbred line names alone. In cases for which classification of alleles at *sh2* was not obvious based on line name alone, lines were designated as ‘unknown’.

### Vitamin TWAS

Using normalized transcript abundance from fresh kernels, we performed TWAS based on read alignments to the B73 and Ia453 reference genomes for 39 existing tocochromanol and carotenoid kernel phenotypes in the same sweet corn diversity panel. Across all traits, 23 unique genes passed a transcriptome-wide FDR threshold of 5% when aligned to B73. Of these, six were located within common support intervals of JL-QTL previously identified with the U.S. maize NAM panel (Diepenbrock et al. 2017; Diepenbrock et al. 2021), including three causal genes: *β-carotene hydroxylase 1* (*crtRB1*), *lycopene ε-cyclase* (*lcyE*), and *vitamin E synthesis4* (*vte4*) (Supplemental Table S5). On a per trait basis, the number of genes passing the significance threshold ranged from zero to nine.

*crtRB1* encodes β-carotene hydroxylase, an enzyme that hydroxylates β-carotene to produce β-cryptoxanthin and zeaxanthin (Yan et al. 2010). In our TWAS, transcript abundance of this gene was significantly associated with four carotenoid traits at a transcriptome-wide level for B73: β-carotene (*P*-value 2.45 × 10^−11^), the ratio of β-carotene to β-cryptoxanthin + zeaxanthin (*P*-value 2.16 × 10^−17^), the ratio of β-carotene to β-cryptoxanthin (*P*-value 3.95 × 10^−14^) and antheraxanthin (*P*-value 3.53 × 10^−6^). *lcyE*, another core carotenoid biosynthesis gene (Harjes et al. 2008), was the top association for α-xanthophylls and the ratio of β-xanthophylls to α-xanthophylls (*P*-value 1.07 × 10^−11^), but only passed the B73 transcriptome-wide 5% FDR threshold for the latter. *vte4*, a core tocochromanol pathway gene (Shintani and DellaPenna 1998), was the top association for five tocopherol traits in this TWAS: αT (*P*-value 3.10 × 10^−11^), γT (*P*-value 3.85 × 10^−8^), γT/(γT +αT) (*P*-value 3.11 × 10^−15^), αT/γT (*P*-value 1.13 × 10^−15^), and δT/αT (*P*-value 3.03 × 10^−8^). Each of these five associations passed the transcriptome-wide 5% FDR threshold for the B73 alignment.

In addition to these *a priori* candidate genes, five other genes were significantly associated with carotenoid traits and 15 genes were significantly associated with tocochromanol traits at the transcriptome-wide level. Of these non-*a priori* candidates, two are transcription factors: *Zea mays MADS18* (Zm00001d010233), which was significantly associated with antheraxanthin (*P*-value 2.33 × 10^−6^), and *HSF transcription factor 11* (Zm00001d034433), which was significantly associated with both γT (*P*-value 1.80 × 10^−5^) and ΣT (*P*-value 2.04 × 10^−5^). Although previous GWAS in this sweet corn diversity panel have identified significant SNPs on the same chromosomes as these genes, neither is physically close with Zm00001d010233 and Zm00001d034433 approximately 33.9 and 8.2 Mb from the nearest significant SNPs, respectively (Baseggio et al. 2019; Baseggio et al. 2020).

Overall, Ia453 TWAS results were similar to B73, with seven genes passing the transcriptome-wide 5% FDR threshold for both alignments (Supplemental Table S6). All three of the causal genes identified with the B73 alignment, *crtRB1, lcyE*, and *vte4*, also passed the significance threshold for Ia453. An additional 11 genes were significantly associated with vitamin traits only in the Ia453 alignment, including two tocotrienol traits (γT3/γT3+αT3, *P*-value 8.4 × 10^−7^; αT3/γT3, *P*-value 1.6 × 10 ^−6^) with *homogentisate geranylgeranyl transferase1* (*hggt1*), a core tocochromanol pathway gene that encodes the first committed step for the synthesis of tocotrienols.

### Pathway-level analysis

The large number of individual tests required for TWAS results in a multiple testing problem that requires correction to control for the Type I error rate, but this can limit the power to detect weaker-effect associations. Using a set of *a priori* candidate genes from the carotenoid and tocochromanol biosynthetic pathways identified in the B73 reference genome (Supplemental Table S4), *P*-values from the B73-aligned TWAS were re-adjusted at a pathway-level. When FDR-adjusted *P*-values were calculated on a pathway-level, eight genes passed the 5% threshold, three for tocochromanol traits and five for carotenoid traits. Five of these genes were located within NAM JL-QTL common support intervals, including *crtRB1, lcyE*, and *vte4*, the three genes identified with the transcriptome-wide FDR threshold (Supplemental Table S7).

Although only significant on the transcriptome-wide level for the Ia453 alignment, transcript abundance of *hggt1* was significantly associated with several tocotrienol traits, γT3/(γT3 +αT3) (*P*-value 9.41 × 10^−6^), αT3/γT3 (*P*-value 1.89 × 10^−5^), and γT3 (*P*-value 1.87 × 10^−4^) on a pathway-level. Several genes involved in the biosynthesis of isoprenoid carotenoid precursors, *diphosphocytidyl methyl erythritol synthase2* (*dmes2*), *deoxy-xylulose synthase2* (*dXs2*), and *cytidine methyl kinase1* (*cmk1*), were identified on a pathway-level for carotenoid-related traits. Transcript abundance of *dmes2* was significantly associated with antheraxanthin and β-xanthophylls, *dxs2* was significantly associated with β-carotene, and *cmk1* was significantly associated with antheraxanthin and the ratio of β-carotene over β-cryptoxanthin + zeaxanthin. In addition to these carotenoid-associated genes, *geranylgeranyl hydrogenase1* (*ggh1*), which is involved in prenyl group and 3,8-divinyl-chlorophyllide biosynthesis I synthesis, was significantly associated with γT/(γT +αT).

### Predictions

A B73-aligned GRM and TRMs from B73 and Ia453 reference genome alignments were used in models with one or multiple relationship matrices for the prediction of all carotenoid and tocochromanol traits. To assess the predictive ability of *a priori* candidate genes from both the carotenoid and tocochromanol pathways, two additional B73-aligned transcriptome-based relationship matrices were evaluated per pathway with one containing only candidate genes (TRM.cand) and the other derived from all remaining non-candidate genes (TRM.non.cand) (Table 1). All models were tested both with and without the inclusion of kernel type mutation (*su1*, *sh2*, or *su1sh2*) as a covariate, but with the exception of some candidate gene-targeted models, its effect on mean predictive ability was less than 5% for all traits and models. Kernel type mutation was included as a covariate for all subsequently reported results. Across the majority of both carotenoid and tocochromanol traits, models incorporating only a GRM (tocochromanols: 0.27 - 0.70; carotenoids: 0.27 - 0.71) outperformed those with a B73-aligned TRM alone (tocochromanols: 0.22 - 0.69; carotenoids: 0.19 - 0.67), with models incorporating both a GRM and TRM (tocochromanols: 0.28 - 0.71; carotenoids: 0.26 - 0.71) obtaining the highest overall predictive abilities. These patterns were consistent for both sets of traits for the Ia453 reference genome.

*A priori* candidate gene-targeted kernels were also tested for predictive ability using B73-aligned transcriptome data. For both the carotenoid and tocochromanol pathways, models containing relationship matrices derived from non-candidate genes alone (TRM.non.cand) outperformed those containing only the candidate gene-targeted TRM (TRM.cand). However, models incorporating a GRM in addition to either of these candidate or non-candidate gene-targeted relationship matrices eliminated this advantage, producing similar predictive abilities with both TRM subset matrices (Supplemental Table S8). In a model including both the TRM.cand and TRM.non.cand relationship matrices (TRM.both), predictive abilities followed similar patterns to models representing the entire transcriptome in a single relationship matrix (TRM.B73), but the mean predictive ability was slightly higher at 0.53 across all carotenoid traits and 0.48 across all tocochromanol traits. The performance difference between TRM.cand models with and without kernel mutation type as a covariate ranged from a 0.17 percentage point difference for ΣT, a trait without endosperm mutation type effects, to a 52.81 percentage point difference for other carotenes, one of the traits with a significant endosperm mutation type effect (Supplemental Table S3).

Although similar patterns were observed across all models for carotenoid traits, mean predictive abilities were slightly higher overall for xanthophylls (0.51 - 0.56) as compared to carotenes (0.48 - 0.51). Lutein, a xanthophyll, had the highest predictive ability amongst carotenoid traits, with predictive abilities ranging from 0.65 (TRM.B73) to 0.71 (GRM and GRM+TRM for both alignments). In contrast, the ratio of zeinoxanthin to lutein had the lowest predictive ability, ranging from 0.19 (TRM.B73) to 0.27 (GRM and GRM+TRM.Ia453). Among models incorporating all transcriptome information for tocochromanol traits, tocotrienol predictive abilities were higher than those for tocopherol traits (Figure 3). The highest tocochromanol trait predictive abilities were obtained for δT3, with predictive abilities ranging from 0.69 (TRM.Ia453) to 0.71 (GRM + TRM for both alignments). γT/(γT+αT) had the lowest predictive abilities overall, ranging from 0.22 (TRM.B73) to 0.29 (GRM + TRM.Ia453).

**Figure 3.**
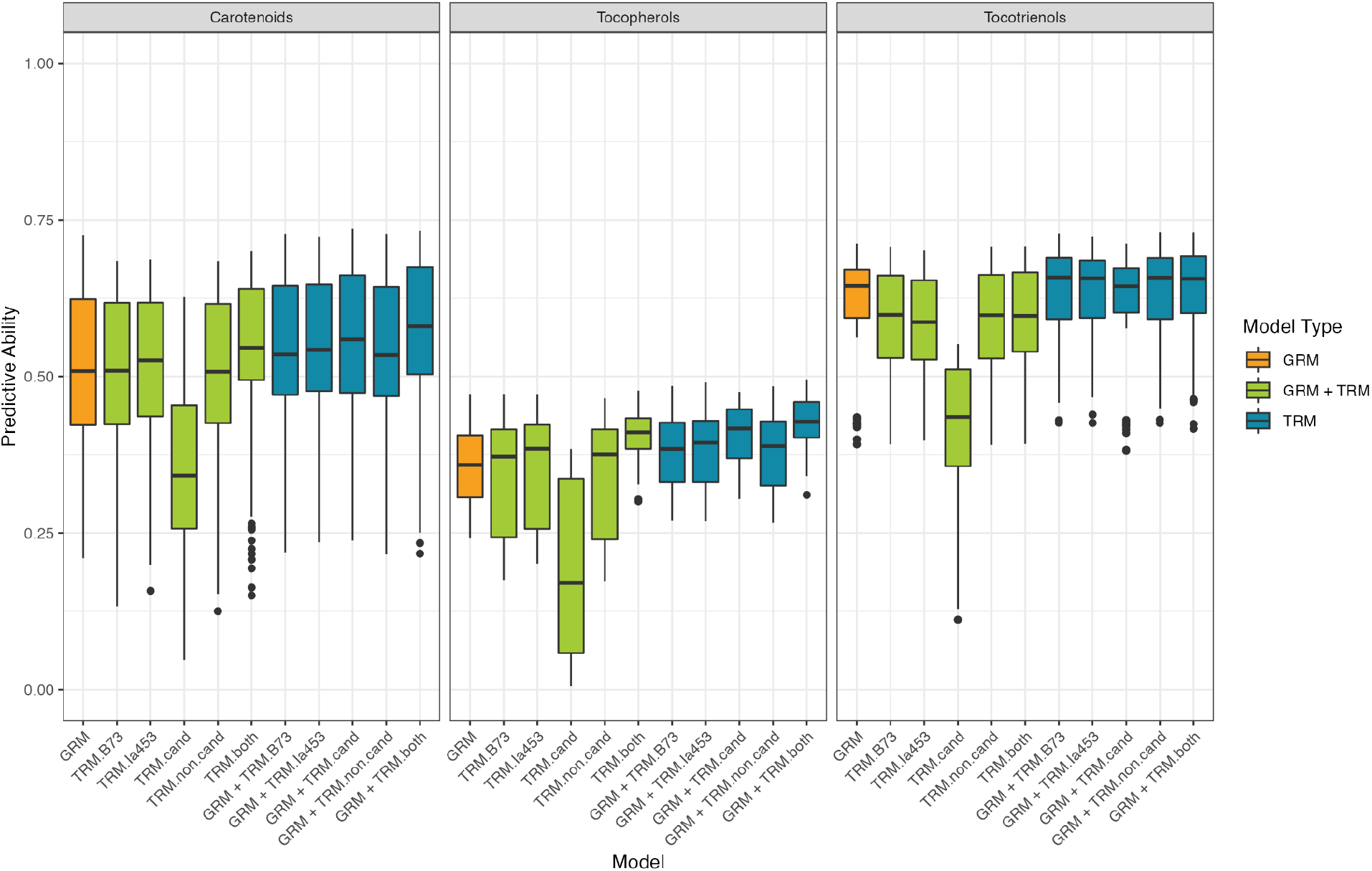
Mean predictive abilities of models incorporating different combinations of relationship matrices. Predictive ability is expressed as the mean Pearson’s correlation between predictions and observed BLUPs for each category of traits across 10 repetitions of five-fold cross-validation. Colors correspond to the types of relationship matrix or matrices included in each model including genomic (GRM), transcriptomic (TRM), or one or more of both types (GRM + TRM).

## DISCUSSION

The biofortification of sweet corn provides an opportunity to address nutritional insufficiencies in populations with high rates of sweet corn consumption. As an intermediate step between genotype and terminal phenotypes, the transcriptome offers valuable insight into mechanisms governing nutrient accumulation in fresh sweet corn kernels that could be exploited for genetic gain by biofortification breeding programs. In this study, we performed transcriptome-wide association and prediction in sweet corn, enabling deeper quantitative genetic analysis of tocochromanol and carotenoid accumulation in fresh kernels.

The sweet kernels characteristic of the sweet corn subpopulation are caused by interruptions to endosperm starch production and storage as a result of mutations in any of eight genes, with *su1* and *sh2* being the most common (reviewed in Tracy et al. 2019). As these genes and the biochemical functions of their encoded enzymes are well-characterized, the kernel mutation type trait represents an ideal case study in which to evaluate the effectiveness of TWAS for causal gene identification in sweet corn.

The *su1* gene encodes an isoamylase, a starch debranching enzyme that is necessary for normal starch biosynthesis in the endosperm (Rahman et al. 1998). Maize kernels with a mutation in this gene have increased levels of simple sugars and phytoglycogen, resulting in a sweet taste and creamy mouth feel (James et al. 1995; Marshall and Tracy 2003). In the progenitors of modern sweet corn in the U.S., the causal variant for *su1* kernel types is a SNP resulting in an amino acid substitution from tryptophan to arginine that renders the enzyme non-functional (Tracy et al. 2006; Kubo et al. 2010). Consistent with the findings of Dinges et al. (2001) and Kubo et al. (2010), transcript abundance of mutant *su1* did not differ from that of wildtype (*Su1*) in our study (Figure 2), whereas a strong association with kernel mutation type was detected through GWAS in this same sweet corn association panel (Baseggio et al. 2019). This is likely because the mutation interferes with protein function without disrupting the intermediate transcriptional process.

While *su1* was the most widely used kernel starch endosperm mutation in commercial sweet corn through the 1960s (Tracy et al. 2019), it has been replaced or accompanied by *sh2* in most modern cultivars. Compared to varieties with mutations in *su1* alone, those homozygous for *shrunken2* mutations have higher levels of simple sugars and a longer shelf life (Garwood et al. 1976; Carey et al. 1982). These compositional changes in mutated kernels are due to interruptions in the activity of ADP-glucose pyrophosphorylase, the enzyme encoded by *sh2*, severely reducing the ability of the kernel to produce starch (Tsai and Nelson 1966). Unlike the single SNP believed to confer the low starch mutation phenotype in *su1*, the most common mutant allele for this gene, *sh2-R*, has major structural rearrangements including an intrachromosomal inversion, retrotransposons, and transposable element insertions (Hu et al.2021; Kramer et al. 2015). These structural events resulted in the splitting of the single progenitor gene into two separate genes, a feature that is reflected in the recently-assembled Ia453 reference genome with the *sh2-R* allele (Hu et al. 2021). Of these two genes, Zm00045a021195 and Zm00045a021196, only the expression of Zm00045a021195 was strongly associated with kernel mutant type in our study (*P*-value 1.32 × 10^−26^). This gene codes for a truncated protein product derived from the last 13 exons compared to the full glucose-1-phosphate adenylyltransferase domain encoded by the functional *Sh2* allele (Hu et al.2021), while the second gene includes only three exons.

Contrary to *sh2-R*, the *sh2-i* mutant allele produces an intermediate phenotype rather than a fully shrunken kernel type (Giroux and Hannah 1994). It is included alongside homozygous mutant *su1* alleles in some commercial sweet corn varieties to both maximize quality and improve seed germination (Dodson-Swenson and Tracy 2015). As inferred from the inbred line names (Supplemental Table S9), this is also the case with a subset of lines in our panel. In agreement with Giroux and Hannah (1994), we found transcript abundance of Zm00001d044129 (the *sh2* locus in B73) in *sh2-i* mutant individuals to approximate the levels of those with wild type *Sh2* alleles (Figure 2). The ability of TWAS to detect these previously-documented subtle differences related to kernel endosperm starch mutation gene expression adds confidence to the potential for the method to be used for accurate characterization of novel associations and biological patterns.

We found TWAS alone to provide a level of detection of known causal genes (*crtRB1, lcyE, vte4*, and *hggt1*) comparable to that of GWAS for fresh kernel carotenoid and tocochromanol traits in this panel (Baseggio et al. 2019; Baseggio et al. 2020). At a genome-wide level, GWAS in sweet corn have identified similar numbers of candidate genes as studies with kernel nutritional quality traits in non-sweet corn maize subpopulations (Lipka et al.2013; Owens et al. 2014; Baseggio et al. 2019; Baseggio et al. 2020). Additional associations were revealed by Lipka et al. (2013) and Owens et al. (2014) through pathway-level analyses, but this type of analysis was not reported by Baseggio et al. (2019; 2020).

Expression of *crtRB1* controls β-carotene concentration in maize endosperm (Yan et al.2010), so it is unsurprising that we found a strong association with it in this study. Previous results have indicated similar associations in this and other maize diversity panels (Harjes et al.2008; Yan et al. 2010; Owens et al. 2014; Suwarno et al. 2015; Azmach et al. 2018; Baseggio et al. 2020). These findings are also supported by the identification of *crtRB1* as a correlated expression and effect QTL (ceeQTL), displaying strong correlations between the JL-QTL allelic effect estimates and expression at multiple stages of kernel development in the U.S. maize NAM panel (Diepenbrock et al. 2021).

Lycopene ε-cyclase adds an ε-ring to one end of lycopene to create α-carotene and has been implicated as a key enzyme at the branch point between α and β carotenoids controlling the accumulation of carotenoids in mature maize kernels (Harjes et al. 2008). The gene encoding this enzyme, *lcyE*, was previously found to be significantly associated with the ratio of β- to α-xanthophylls in this sweet corn panel (Baseggio et al. 2020) and with six carotenoid traits in the Goodman-Buckler panel (Owens et al. 2014). Though this ratio trait was not tested in the U.S. maize NAM panel, *lcyE* falls within the QTL region identified by JL analysis for six other carotenoid traits (Diepenbrock et al. 2021). *lcyE* was also designated by Diepenbrock et al. (2021) as a ceeQTL Given the major effects observed in these previous studies, it is confirmatory that we were also able to detect strong associations between *lcyE* and several carotenoid traits through TWAS.

While *dxs2* has previously been associated with carotenoid content in GWAS and ceeQTL analysis of the U.S. maize NAM and Goodman-Buckler panels (Owens et al. 2014;Diepenbrock et al. 2021), significant associations of *dmes2* and *cmk1* with carotenoid-related traits have not previously been reported in maize despite assignment as *a priori* candidates. *dxs2* encodes a paralog of 1-deoxy-D-xylulose 5-phosphate synthase (DXS), an enzyme involved in IPP synthesis upstream of both the carotenoid and tocochromanol biosynthetic pathways. It was first associated with maize grain carotenoids by Owens et al. (2014) at a pathway-level and has since been identified as a JL-QTL and ceeQTL in the U.S. maize NAM panel for both carotenoid and tocochromanol traits (Diepenbrock et al. 2017; Diepenbrock et al. 2021). *dxs2* was associated with transcript abundance in a pathway-level TWAS of β-carotene but not for any tocochromanol traits nor through GWAS of traits from either trait group in this panel (Baseggio et al. 2020; Baseggio et al. 2019). The PVE for *dxs2* in the US maize NAM panel was 8.1% for β-carotene, larger than the PVEs for the same gene when associated with tocotrienol traits (2.5-5.7% PVE), potentially explaining the difference in our ability to detect an association in the sweet corn association panel.

For both tocopherols and tocotrienols, the protein encoded by *vte4* methylates γ- and δ-species to form α- and β-species, respectively. At least one known allele of *vte4* has been shown to affect transcript abundance of the gene (Li et al. 2012), with further support of expression-mediated phenotypic control by the identification of this gene as a ceeQTL in the U.S. NAM panel (Diepenbrock et al. 2017). Although associations have been found between tocopherols and *vte4* in both GWAS (Baseggio et al. 2019) and TWAS (this study) in our sweet corn panel, no significant associations have been identified between tocotrienol traits and this gene in sweet corn association studies (Baseggio et al. 2019) despite detected associations at the genome-wide (Diepenbrock et al. 2017) and pathway-level (Lipka et al. 2013) in non-sweet corn maize inbred panels. While there is not a clear biological reason for this finding, the lack of signal present for tocotrienols in GWAS and TWAS in this panel suggests that unlike founders of the maize U.S. NAM panel, the sweet corn panel may contain alleles in which *vte4* is more poorly expressed in the endosperm, the major site of tocotrienol synthesis in kernels (Grams et al. 1970), which would explain the association with tocopherols (whose major site of synthesis is embryo; Grams et al. 1970) and absence of strong association with tocotrienols.

While *hggt1* did not pass the transcriptome-wide 5% FDR threshold for B73-aligned transcripts in this study, significant associations were identified between this gene and several tocotrienol traits at a transcriptome-wide level for the Ia453 alignment. Additional significant associations with *hggt1* were identified at a pathway-level for B73. This gene has previously been found to be highly expressed in the endosperm (Stelpflug et al. 2016), the primary location for tocotrienol synthesis (Grams et al. 1970), during maize kernel development (Diepenbrock et al. 2017). Markers in strong LD with *hggt1* were previously found to be significantly associated with tocotrienol traits in this and other maize diversity panels (Baseggio et al. 2019; Lipka et al.2013). These associations are further supported by Diepenbrock et al. (2017), who found *hggt1* to be a ceeQTL in addition to it explaining a large percentage of phenotypic variance for three tocotrienol traits in the U.S. maize NAM panel. In contrast, we did not detect an association of *vitamin E synthesis1* (*vte1*) with tocotrienols at either the transcriptome-wide or pathway-level, although strong associations between *vte1* and tocotrienols have been detected via GWAS in this sweet corn panel (Baseggio et al. 2019). While it is possible that the peak expression of *vte1* alleles occurs outside of the RNA sampling time point, it is more likely that the causal variant(s) does not have a strong expression effect.

Although the *protochlorophyllide reductase* (*por*) homologs *por1* and *por2* together account for the largest PVEs observed for total tocopherols in the U.S. NAM panel (Diepenbrock et al. 2017), neither designated ceeQTL was significantly associated with tocochromanol traits in this study.*por1* was not expressed in the majority of our samples, thus it did not pass necessary quality filtering thresholds for inclusion in this study. In contrast, *por2* was included in TWAS but was not amongst the most significant associations for any trait in this study. Severe bottlenecks during sweet corn population development (Tracy 1997; Whitt et al. 2002) may have resulted in large-effect causal variants becoming rarer or small-effect causal variants increasing to higher frequency, thus limiting the detection of either *por* gene in a panel consisting exclusively of sweet corn lines.

### Predictions

For many traits, including αT3, δT3, δT, γT, γT3, ΣT, ΣT3, ΣT3 +ΣT, antheraxanthin, β-cryptoxanthin, β-xanthophylls, and zeaxanthin, we observed substantial increases in the predictive ability of models incorporating both a GRM and TRM over those with only a GRM, regardless of the reference genome to which transcripts were aligned. This indicates that the relative importance of including transcript abundance data varies depending on the predicted trait, consistent with previous findings for other maize traits (Guo et al. 2016; Schrag et al.2018). Furthermore, in concordance with the GRM-only predictive abilities obtained by Baseggio et al. (2019; 2020), GRM + TRM, GRM-only and TRM-only accuracies were lower for tocopherol traits as compared to tocotrienols and carotenoids for all reference genome alignments.

Due to strong associations of tocotrienol and some carotenoid traits with kernel mutant type in this sweet corn association panel (Baseggio et al. 2019; Baseggio et al. 2020), all models were tested both with and without the inclusion of kernel type mutation (*su1*, *sh2*, or *su1sh2*) as a covariate. There was no clear pattern of improvement with or without kernel type across models incorporating genome-wide or transcriptome-wide relationship matrices in either trait set, likely because population structure, which is highly related to kernel mutant type in this panel (Baseggio et al. 2019), is represented in the genomic and transcriptomic relationship matrices themselves. However, for both carotenoid and tocochromanol traits previously identified as having a significant endosperm mutation type effect in this panel (Baseggio et al. 2019; Baseggio et al. 2020), the inclusion of kernel mutant type as a covariate in models with only a TRM.cand boosted model performance dramatically. A similar pattern was also observed by Baseggio et al. (2019; 2020) for pathway-level and QTL-targeted genomic prediction models for these traits.

As was the case with pathway-level and QTL-targeted marker datasets (Baseggio et al. 2019; Baseggio et al. 2020), *a priori* candidate gene transcript abundance datasets alone did not predict the majority of carotenoid or tocochromanol traits as well as genome- or transcriptome-wide markers in this panel, regardless of the inclusion of kernel mutant type as a covariate. Prediction models incorporating only TRM.cand outperformed those with TRM.non.cand alone for only six traits: β-carotene, the ratio of β-carotene over β-cryptoxanthin + zeaxanthin, the ratio of β-xanthophylls over α-xanthophylls, αT, γT/(γT+αT), and αT/γT. This gap in performance was minimized by the addition of genome-wide marker data, suggesting that missing information in the transcriptome subset relationship matrices is made up for by the presence of genomic markers.

## CONCLUSION

Leveraging a high quality transcriptomic dataset collected from fresh kernels, this study represents the first formal TWAS and transcriptomic prediction in sweet corn. As demonstrated through both kernel mutation type and vitamin traits, TWAS is a viable approach for the detection of causal genes in this system. On a transcriptome-wide level, TWAS enabled the identification of four genes known to control vitamin accumulation: *crtRB1, lcyE, vte4*, and *hggt1*. Pathway-level analysis further facilitated the detection of *dxs2* and three genes that have not previously been reported in maize association studies: *dmes2*, *cmk1*, and *ggh1*. The use of transcriptome-wide data increased predictive ability when incorporated into models alongside genome-wide markers, but the relative boost was trait dependent. Overall, predictive abilities ranged from moderate to high, but *a priori* candidate gene-targeted prediction models underperformed compared to those using transcriptome-wide datasets.

Given that this study explored transcriptomic data from a single environment and time point, the collection and analysis of kernel RNA at additional developmental stages or of other endophenotypes may help to more fully understand the biological mechanisms underpinning vitamin accumulation in fresh sweet corn kernels. These data could further be incorporated into prediction models to improve predictive abilities for these and other kernel traits. Taken together, this work contributes to a better understanding of the role of gene expression in the accumulation of carotenoids and tocochromanols in fresh sweet corn kernels, an additional step towards the biofortification of sweet corn for increased nutritional quality.

## Author contributions

JH: Funding acquisition, Conceptualization, Investigation, Formal analysis, Writing - Original Draft, Writing - Review & Editing

RT: Formal analysis, Writing - Review & Editing

JCW: Formal analysis, Writing - Review & Editing

NK: Investigation

DW: Formal analysis, Writing - Review & Editing

JPH: Formal analysis

DD: Funding acquisition, Resources, Writing - Review & Editing

CRB: Funding acquisition, Resources, Supervision, Writing - Review & Editing

MAG: Funding acquisition, Conceptualization, Supervision, Resources, Writing - Review & Editing

## Abbreviations

αT: α-Tocopherol
αT3: α-Tocotrienol
γT: γ-Tocopherol
γT3: γ-Tocotrienol
δT: δ-Tocopherol
δT3: δ-Tocotrienol
*aeduwx*: *amylose-extender:dull:waxy*
BLUP: best linear unbiased predictor
*bt2*: *brittle2*
*cmk1*: *cytidine methyl kinase1*
*crtRB1*: ß-*carotene hydroxylase*
*dmes2*: *diphosphocytidyl methyl erythritol synthase2*
*dxs2*: **deoxy-xylulose synthase2**
FDR: false discovery rate
GBS: genotyping-by-sequencing
GBLUP: genomic best linear unbiased prediction
*ggh1*: *geranylgeranyl hydrogenase1*
GRM: genomic relationship matrix
GWAS: genome-wide association study
*hggt1*: *homogentisate geranylgeranyltransferase1*
HPLC: high-performance liquid chromatography
JL: joint-linkage
*lcyE*: *lycopene epsilon-cyclase*
LD: linkage disequilibrium
MEP: methylerythritol phosphate
MLMM: multi-locus mixed-model
NAM: nested association mapping
QTL: quantitative trait locus
RDA: recommended daily allowance
*sh2*: *shrunken2*
SNP: single-nucleotide polymorphism
*se1*: *sugary enhancer1*
*su1*: *sugary1*
TRM: transcriptomic relationship matrix
ΣT: total tocopherols
ΣT3: total tocotrienols
ΣT3 + ΣT: total tocochromanols
*vte1*: *tocopherol cyclase*
*vte4*: *γ-tocopherol methyltransferase*

## DATA AVAILABILITY

All raw 3’ mRNA-seq data are available from the NCBI Sequence Read Archive under BioProject xxx (to be released upon publication). Code is available from GitHub at https://github.com/GoreLab/SweetCornRNA.

## ACKNOWLEDGEMENTS

This research was supported by the National Institute of Food and Agriculture; the USDA Hatch under grant numbers 100397, 1010428, 1013637, 1013641, and 1023660 (M.A.G.) and AFRI EWD Predoctoral Fellowship 2019-67011-29606 (J.H.); the National Science Foundation (IOS-1546657 to D.D.P, C.R.B, and M.A.G.); Cornell University startup funds (M.A.G.). We thank Matt Baseggio and other current and past members of the Gore lab for their efforts in pollination, harvest, and sample preparation.

## SUPPLEMENTAL MATERIAL

- **Supplemental Table S1.** Sample and gene counts throughout the normalized transcript abundance quality control pipeline
- **Supplemental Table S2.** Line, gene, and marker counts in analysis pipelines
- **Supplemental Table S3.** Optimal transcriptome-wide association study models as selected by Bayesian information criterion
- **Supplemental Table S4.** List of a priori candidate genes included in candidate gene-targeted transcriptomic prediction matrices
- **Supplemental Table S5.** Vitamin trait significant associations at the transcriptome-wide level for the B73 v4 reference genome
- **Supplemental Table S6.** Vitamin trait significant associations at the transcriptome-wide level for the Ia453 reference genome
- **Supplemental Table S7.** Vitamin trait significant associations at the pathway-level for the B73 v4 reference genome
- **Supplemental Table S8.** Genomic and transcriptomic predictive abilities for all models
- **Supplemental Table S9.** Endosperm mutation assignments

## CONFLICT OF INTEREST

The authors declare no conflict of interest.

## Notes

### Competing Interest Statement

The authors have declared no competing interest.

